# Prevalence of parasitic infections among recent immigrants to Chicago

**DOI:** 10.1101/575779

**Authors:** Jesica A. Herrick, Monica Nordstrom, Patrick Maloney, Miguel Rodriguez, Kevin Naceanceno, Gloria Gallo Enamorado, Rojelio Mejia, Ron Hershow

## Abstract

**Background:** Parasitic infections are likely under-recognized among immigrant populations in the United States (US). We conducted a cross-sectional study to evaluate the frequency of such infections among recent immigrants in Chicago and to identify predictive factors for parasitic infections.

**Methodology and principal findings:** 133 recent immigrants were enrolled, filling out a standardized questionnaire regarding medical history and exposures and providing blood and stool samples for evaluation. Fifteen of 125 subjects (12%) who provided a blood or stool sample for testing were found to have evidence of current or prior infection with a pathogenic parasite, of which *Toxocara* spp. (8 subjects, 6.4%) and *Strongyloides stercoralis* (5 subjects, 4%) were most commonly identified. Parasitic infection was more likely among subjects who had immigrated within the previous 2 years and those with a self-reported history of ever having seen worms in the stool. Infected individuals were likely to have multiple nonspecific physical complaints; however, classic symptoms of parasitic infections (skin rashes, diarrhea, *etc.)* were not increased among infected individuals. The most useful surrogate markers identified for parasitic infections were an elevated Immunoglobulin E level (seen in 7/15 subjects with parasitic infections, 46.7% and 22/110 uninfected individuals, 20%, p=0.04) and the presence of *Blastocystis hominis* cysts on Ova & Parasite exam (detected in 5/13 subjects with parasitic infections who provided a stool sample, 38.5% and 5/98 uninfected subjects, 5.1%, p=0.002). In contrast, the Absolute Eosinophil Count (typically thought of as an indicator of parasites) was not found to be a good screening test for parasitic infections in this study.

**Conclusions:** Our study found that parasitic infections are common in recent US immigrants, which highlights an important health disparity among a vulnerable population. Further, we found that classically used symptoms and laboratory tests had a low predictive value for parasitic infections in this population.

**AUTHOR SUMMARY:** Parasitic infections, though rare in the United States (US), are common in many areas of the world including the regions of origin of many US immigrants. However, the prevalence rates and health impacts of these infections in immigrant populations are undefined. We conducted a study to identify the frequency of parasitic infections among healthy immigrants in one community, recruiting 133 immigrants from 28 countries. Subjects completed a standardized questionnaire regarding symptoms and infection risk-factors and provided blood and stool samples for testing. Twelve percent of subjects in our study had evidence of current or previous pathogenic parasitic infections. Symptoms and risk factors classically thought to be associated with parasitic infection (allergic symptoms, elevated blood eosinophil counts, *etc.)* were common among enrolled subjects, but did not differ significantly between those with and without evidence for infection. Overall, our results suggest that many immigrants, even those who are asymptomatic, may have undiagnosed parasitic infections. These results highlight an important health disparity among a vulnerable underserved population in the US. As most of these infections are easily treatable, more research should be done to further characterize the optimal testing strategies for recent immigrants.

## INTRODUCTION

In some regions and among select populations, parasitic diseases are believed to be a significant and under-recognized public health problem within the United States (US).[1–7] However, little is known about the current epidemiology of these infections.[3, 8] Although exact numbers are not available, studies have suggested that within the US there may be up to 300,000 people with Chagas disease,[9–15] 1.3 – 2.8 million with serological evidence of exposure to *Toxocara* spp.,[3, 16] 4 million with soiltransmitted helminths,[3, 17, 18] 1.2 million with giardiasis,[19] 41,400 – 169,000 with cysticercosis,[3] and approximately 8,000 with schistosomiasis.[3] These infections represent an important health disparity in the US as higher infection rates for some parasites are believed to occur among those living in poverty in underserved rural and inner-city areas.[2, 3, 20–23]

Parasitic infections are believed to be an especially significant health problem within US immigrant populations. Many immigrants come from areas where some parasites are endemic.[3, 24–27] Additionally, recent immigrants may be more likely to have a decreased socioeconomic status and therefore frequently reside in the underserved areas within the US that have an increased prevalence of parasitic infections.[2–4, 22] This population thus possesses a dual risk for parasites.[7] However, published prevalence estimates vary widely and therefore the true prevalence and health impacts of parasitic infections on US immigrant populations is unknown.

Both diagnostic and methodologic limitations to research may explain the lack of information regarding parasitic infections among US immigrants. Diagnostic inaccuracies can occur because these infections frequently have nonspecific symptoms or are asymptomatic, and laboratory tests for parasites can have significant limitations. For example, many of the current diagnostic tests for parasitic infections have poor sensitivity. Additionally, these tests are frequently unavailable in the community.[4, 6, 7, 17, 25, 28, 29] Serologic testing, the primary method of diagnosis for many parasitic infections, has several important shortcomings, which contributes to the lack of accurate information regarding these infections. These limitations include: an inability to distinguish between active and past infections, the presence of significant crossreactivity between serologic tests for different parasites, and the fact that some infections (such as many stool helminthiases) do not cause a detectable serologic response.[30–32]

Methodologic challenges to research studies may also partly account for the wide variations in reported prevalence estimates of parasitic infections.[1, 3, 4] For example, past estimates of Chagas disease prevalence were based on testing of blood-donors. This method likely underestimated the number of infected individuals due to a “healthy-donor” effect and due to the removal of those with previous positive tests from the pool of potential blood donors.[33] A second commonly used method to obtain prevalence estimates involves multiplying the proportion of people infected with a specific disease within a country by the number of immigrants from that country living in the US. However, within endemic countries there can be large differences in infection risk across different populations and infection risks among those who emigrated may be different from that of the general population within a given country.[10, 11, 24, 34–36]

Given the limited data regarding the prevalence of parasitic infections, it is unsurprising that the health impacts imposed by parasites on US immigrant populations are poorly understood. Many of the published studies evaluating health impacts of parasitic infections on immigrants either screened for only a small number of parasites or screened immigrants from a single country, and many were retrospective.[9, 11, 13, 14, 25, 26, 37, 38] Improved data regarding the prevalence and impact of these infections are badly needed as parasitic infections can have serious health consequences.[2, 3, 5–7, 17, 20, 39] For example, Chagas disease causes cardiomyopathy and/or esophageal or colonic dilation in up to 20-30% of those infected;[9, 40] those with Toxocariasis may have an increased risk for asthma;[41–44] and intestinal parasite infection has been associated with delayed cognitive development and impaired nutrition in some studies.[45, 46] Parasitic infections have also been shown to increase the severity of illnesses due to other infectious diseases (such as tuberculosis and HIV).[1]

Most studies evaluating the impact of parasitic infections in the US have focused on the Southern states because of a warmer climate (facilitating the acquisition of soil-transmitted helminths) and a comparatively high population of immigrants.[5, 12, 13, 22, 47] According to US Census reports, however, Illinois is among the states with the highest percentages of immigrants; 10-15% of the state’s population is foreign born.[48] Thus, this study was conducted with the overall aim of evaluating the prevalence of parasitic infections among recent immigrants living in Chicago. As many of the symptoms of parasitic infections are nonspecific, an additional objective of our study was to identify any symptoms, signs, or laboratory tests that could be used in this population as predictors of the presence of parasites.

## METHODS

### Ethics statement

The study was approved by the Institutional Review Board of the University of Illinois at Chicago and written informed consent was obtained from all participants. Adult subjects signed a consent form, and for enrolled individuals between the ages of 10 and 18, the subject signed an assent form and a parent or guardian signed a parental permission form. For all study activities, study procedures were explained to the participant in their stated language of preference (using an interpreter if needed). Each participant received $40 to compensate them for their time. Treatment for parasitic infections was not a part of the study, but all subjects were notified of their results and those with positive tests were offered the possibility of seeing one of the study authors (JH) in clinic to discuss treatment options.

### Study design

Patients were enrolled for this cross-sectional study between November 2014 and July 2016. Subjects were recruited through collaboration with local organizations that provide services to immigrants as well as at English as a Second Language classes, the Mexican consulate in Chicago, through a mass email sent to students enrolled at the University of Illinois at Chicago, and at community health fairs. Subjects were also recruited from community health clinics as long as they were accessing the health care system for reasons clearly unrelated to a parasitic infection (e.g., patients presenting for routine physical exams). Subjects were eligible for inclusion if they were: ages 10-80; willing to undergo a blood draw and provide a stool sample; and an immigrant living in the US mainland for less than 5 years and were originally from Africa, Asia, South America, Central America, Mexico, the Caribbean islands, or Puerto Rico. Although they are U.S. citizens and not immigrants from non-US territories, Puerto Ricans were specifically included in this study because previous research suggested that parasitic infections were common among the Puerto Rican population in Chicago.[49] The five-year time point for living in the US was chosen because immigrants have been shown to have the highest risk for parasitosis in the first 5 years after emigration[49] and based on the average expected lifespan of hookworms, *Ascaris lumbricoides* and *Trichuris trichiura*, a primary focus of our investigations.[50] Subjects were excluded from the study if they: 1) were currently pregnant, as the immunosuppression of pregnancy may cause a different presentation of parasitic infection and one of the aims of the study was to identify factors predictive of parasitosis; or 2) they had taken antiparasitic medications since moving to the continental US.

Upon enrollment, a standardized questionnaire (Supplemental Appendix) was administered to collect information regarding demographics, risk factors for parasitic infections, and symptoms. Participants then provided clinical samples (blood and stool) to be tested for evidence of parasitic infections as described below.

### Laboratory testing

Whole blood for a Complete Blood Count was collected in an Ethylenediaminetetraacetic acid (EDTA) tube and processed in the clinical laboratory at the University of Illinois at Chicago (BXH 800 or BXH 1600, Beckman Coulter, Brea, CA) within 12 hours. Serum samples were collected in a serum separator tube (SST) and centrifuged within 1 hour of collection. A portion of the supernatant was sent for measurement of Immunoglobulin E (IgE) level (Quantitative ImmunoCAP^®^ Fluorescent Enzyme Immunoassay, Arup Laboratories, Salt Lake City, UT). The remainder of the supernatant was stored at −80 °C until samples were shipped as a batch to the Centers for Disease Control and Prevention, where serologic testing was performed for infections endemic to each subject’s country of origin (Supplemental Table 1).

Antibody responses to cysticercosis, strongyloidiasis, and toxocariasis were determined using a multiplex bead-based assay (MBA).[51–53] For cysticercosis, 3 recombinant antigens (rGP50, rT24H, and sTs18var1) were used; one recombinant antigen was used for both strongyloidiasis (Ss-NIE-1) and toxocariasis (rTc-CTL-1). Briefly, the antigens were coupled to carboxylated magnetic particles, and then incubated with sera (diluted 1:100 in PBS/0.3% Tween-20/5% milk) for 30 minutes. After two washes with PBS/0.3% Tween-20, the complex was detected by a biotinylated conjugate which then reacted with streptavidin-phycoerythrin. The mean fluorescence intensity was measured using the MagPix instrument (Luminex corporation, Austin, TX). Positivity for each antibody response was determined using specific cut-off points for each antigen (which had been previously defined[51–53]).

Antibody against *T. cruzi* antigens was detected using a commercial kit (Weiner Lab, Argentina) and following the company protocol instructions.[54–56] After adding serum, the plate was incubated for 30 min while shaking at room temperature. After the conjugate was added, the reaction ran for an additional 30 minutes. Substrate (TMB) was added and color development was stopped by adding 1N H_2_SO4 according to the kit instructions[55]. The optical density at 450nm was detected using the Versamax system.

All serum specimens were initially tested by FAST-ELISA using *Schistosoma mansoni* adult microsomal antigen and then by a species-specific immunoblot appropriate to the subject’s country of origin. These procedures and the methods used to interpret results have been discussed in previous publications.[57–59]

Patients also provided a stool sample for standard Ova & Parasite (O&P) direct microscopy (Arup Labs Qualitative Concentration/Trichrome Stain/Microscopy, Salt Lake City, UT), and a multi-parallel quantitative polymerase chain reaction (qPCR) which tests for eight common gastrointestinal pathogens *(Ancylostoma, Ascaris, Cryptosporidium, Entamoeba, Giardia, Necator, Strongyloides*, and *Trichuris).* Only one stool sample was requested for O&P due to our desire to compare O&P directly with qPCR and because the yield of collecting multiple samples for O&P exam has been shown to be low in immigrant populations.[60] The stool samples for qPCR were frozen without fixatives at −80 °C and shipped as a batch to the Baylor College of Medicine where DNA was extracted and the qPCR was conducted according to previously published methods.[61–63]

### Statistical analysis

Statistical analyses were performed using Prism 7 (GraphPad, San Francisco, CA) and SAS (Cary, NC). Data were summarized by using frequency with percentage for categorical variables. For continuous variables, geometric mean with standard deviation or median with range or interquartile range were used (medians were used as measures of central tendency for highly skewed variables). For categorical variables, prevalence ratios (PR) were used to evaluate relationships between covariates and the main outcome (the presence of a parasitic infection) and Fisher’s Exact test or Chi-squared test were used to determine significance. Mann-Whitney U test was used to evaluate for associations between continuous variables and the presence of a parasite. Correlations were calculated using Spearman’s rank correlation coefficient. Kappa values were used to compare results from stool O&P exam to qPCR testing.

## RESULTS

### Patient characteristics

Seven hundred and thirty-eight people were approached about the study, and 133 were enrolled (Supplemental Figure 1). All enrolled subjects filled out the study questionnaire (unedited results shown in Supplemental Table 2); 125 (94%) of enrolled subjects provided blood and 113 (85%) provided stool samples to be tested for parasitic infections (however, for 2 of the 113 subjects, only qPCR could be performed as they did not return samples for O&P exam). The mean age of enrolled subjects was 32 years, and 60/133 (45.1%) were male (Table 1). Participants came from 28 different countries. Most subjects (95/133, 71.4%) reported an annual household income of less than $20,000 per year. As shown in Table 1, few subjects (25/133, 18.8%) had been raised in a rural environment. Despite this, many subjects reported having been exposed to animals (65/133, 48.9%), well-water (65/133, 48.9%), and bathing in ponds or streams (58/133, 43.6%) in their country of origin.

**Table 1.** Demographics, symptoms, and laboratory results of 133 enrolled subjects

| Characteristic | Results<sup>a</sup> |
| --- | --- |
| Age, mean years (standard deviation) | 32 (17.3) |
| Gender, male:female, N (% male) | 60:73 (45.1) |
| Country/region of origin, N (%) |  |
| Asia (excluding India) | 33 (24.8) |
| Mexico | 30 (22.6) |
| India | 26 (19.5) |
| Central/South America | 22 (16.5) |
| Africa | 13 (9.8) |
| Middle East | 5 (3.8) |
| US (Puerto Rico) | 4 (3) |
| Mean number of years in school (standard deviation) | 9.9 (5.2) |
| Annual household income in US dollars, N (%) |  |
| < 20,000 | 95 (71.4) |
| 20,000 - 40,000 | 30 (22.6) |
| 40,000 - 60,000 | 5 (3.8) |
| 60,000 - 100,000 | 2 (1.5) |
| > 100,000 | 1 (0.8) |
| Length of time living in the continental United States<sup>b</sup>, mean years<br>(range, standard deviation in years) | 2 (10 days–5<br>years, 1.6) |
| Raised in rural environment, N (%) | 25 (18.8) |
| Exposure history prior to immigration, number yes (%) |  |
| Frequently walked barefoot | 76 (57.1) |
| Close contact with animals | 65 (48.9) |
| Used a well as source of drinking water | 65 (48.9) |
| Bathed in streams or ponds | 58 (43.6) |
| Lived in a house with a thatched roof | 25 (18.8) |
| Treatment with antiparasitic medication prior to emigration, number yes (%) | 46 (34.6) |
| Symptoms reported by the patient on the day of study enrollment<sup>c</sup>, N (%) |  |
| Gastrointestinal | 32 (24.1) |
| Constitutional | 25 (18.8) |
| Musculoskeletal | 25 (18.8) |
| Dermatologic | 13 (9.8) |
| Pulmonary | 12 (9) |
| Cardiovascular | 10 (7.5) |
| Allergic | 3 (2.3) |
| Immunoglobulin E (IgE) results |  |
| Subjects with elevated Immunoglobulin E, N (%) | 29/125<sup>d</sup> (23.2) |
| Immunoglobulin E level, IU/mL, median (range, standard deviation) | 57 (3-7732, 706) |
| Absolute eosinophil count (AEC) results |  |
| Subjects with elevated absolute eosinophil count, N (%) | 12/123<sup>e</sup> (9.8) |
| Absolute eosinophil count, median cells/microliter<br>(range, standard deviation) | 200 (0-1600,<br>240) |
| Pathogenic parasitic infection, N (%) | 15/125<sup>f</sup> (12) |

### Laboratory testing

The median IgE among the 125 subjects from whom blood samples were obtained was 57 IU/mL (range 3-7732); 29 (23.2%) subjects had elevated IgE levels. Twelve subjects (9.8%) were found to have eosinophilia, and the median absolute eosinophil count (AEC) level was 200 cells/UL (range 0-1600 cells/UL, Table 1). There was a weak correlation between IgE and AEC levels in adults (r=0.23, p=0.02; children were excluded from this correlation as they have different normal ranges for IgE).

### Parasitic infections

Fifteen of 125 subjects (12%) who provided a clinical sample for testing were found to have evidence of current or prior infection with a pathogenic parasite species. As shown in Table 2, the most common infections identified were *Toxocara* spp., seen in 8 subjects (6.4%), followed by *Strongyloides stercoralis* (5 subjects, 4%). Of subjects found to have evidence of parasitic infection, 3 subjects (3/15, 20%) had positive tests for more than one pathogenic parasitic species (Table 2).

**Table 2.**
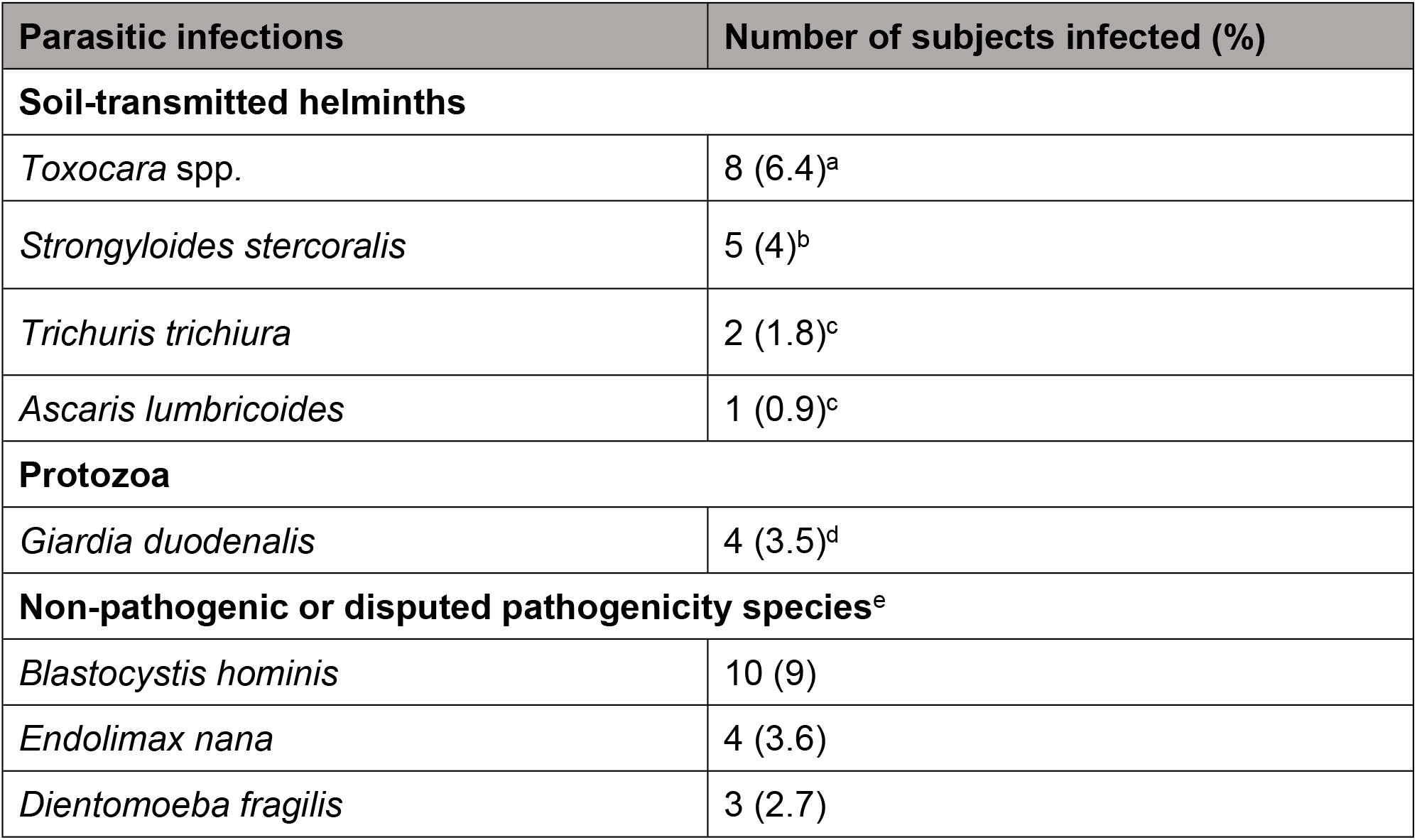
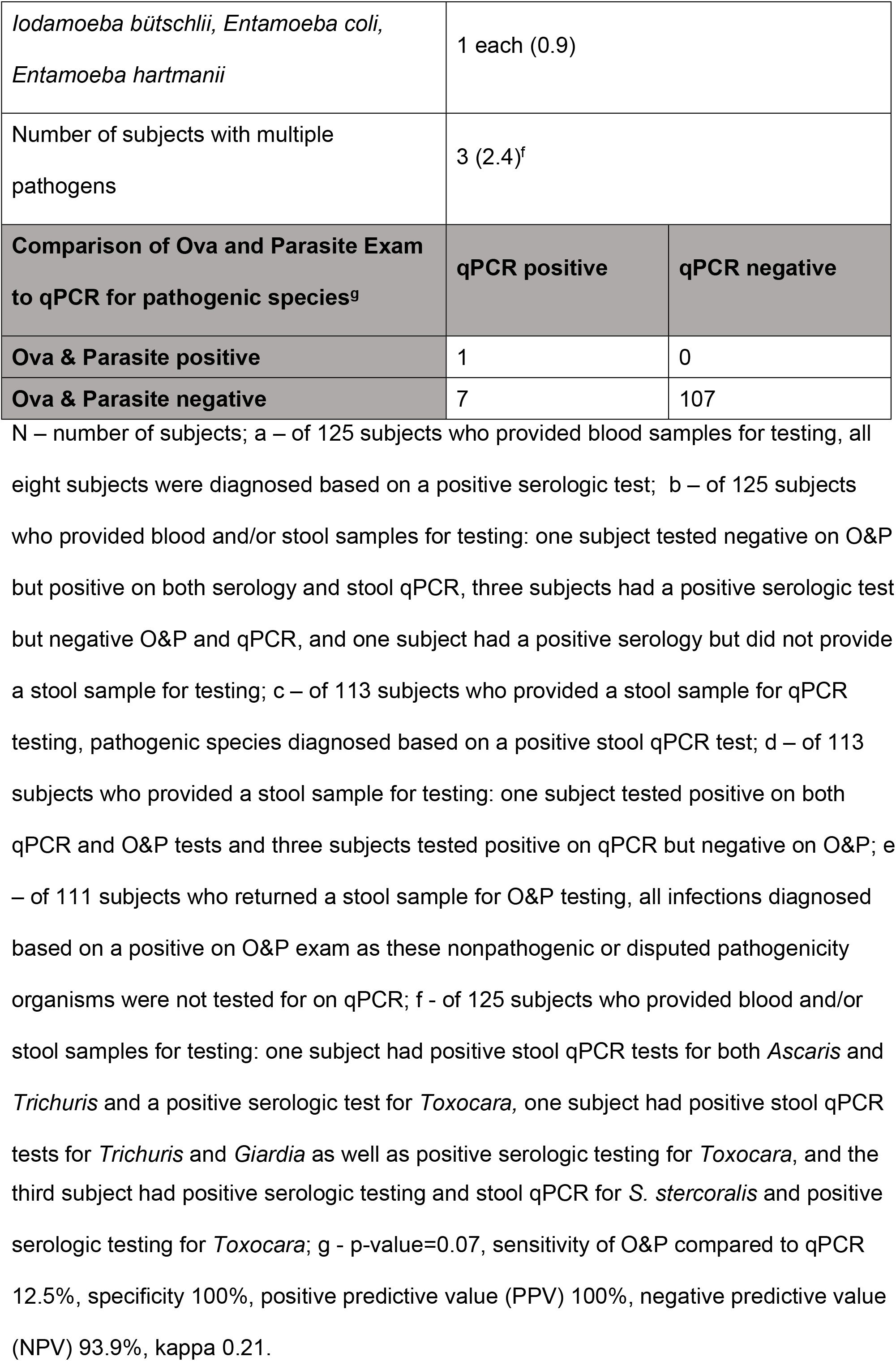
Parasitic infections diagnosed among recent immigrants living in Chicago

| Parasitic infections | Number of subjects infected (%) |
| --- | --- |
| **Soil-transmitted helminths** |  |
| *Toxocara* spp. | 8 (6.4)<sup>a</sup> |
| *Strongyloides stercoralis* | 5 (4)<sup>b</sup> |
| *Trichuris trichiura* | 2 (1.8)<sup>c</sup> |
| *Ascaris lumbricoides* | 1 (0.9)<sup>c</sup> |
| **Protozoa** |  |
| *Giardia duodenalis* | 4 (3.5)<sup>d</sup> |
| **Non-pathogenic or disputed pathogenicity species<sup>e</sup>** |  |
| *Blastocystis hominis* | 10 (9) |
| *Endolimax nana* | 4 (3.6) |
| *Dientamoeba fragilis* | 3 (2.7) |

|  |  |  |
| --- | --- | --- |
| *Iodamoeba bütschlii*, *Entamoeba coli*,<br>*Entamoeba hartmanii* | 1 each (0.9) |  |
| Number of subjects with multiple pathogens | 3 (2.4)<sup>f</sup> |  |
| **Comparison of Ova and Parasite Exam to qPCR for pathogenic species<sup>g</sup>** | **qPCR positive** | **qPCR negative** |
| **Ova & Parasite positive** | 1 | 0 |
| **Ova & Parasite negative** | 7 | 107 |

N – number of subjects; a – of 125 subjects who provided blood samples for testing, all eight subjects were diagnosed based on a positive serologic test; b – of 125 subjects who provided blood and/or stool samples for testing: one subject tested negative on O&P but positive on both serology and stool qPCR, three subjects had a positive serologic test but negative O&P and qPCR, and one subject had a positive serology but did not provide a stool sample for testing; c – of 113 subjects who provided a stool sample for qPCR testing, pathogenic species diagnosed based on a positive stool qPCR test; d – of 113 subjects who provided a stool sample for testing: one subject tested positive on both qPCR and O&P tests and three subjects tested positive on qPCR but negative on O&P; e – of 111 subjects who returned a stool sample for O&P testing, all infections diagnosed based on a positive on O&P exam as these nonpathogenic or disputed pathogenicity organisms were not tested for on qPCR; f - of 125 subjects who provided blood and/or stool samples for testing: one subject had positive stool qPCR tests for both *Ascaris* and *Trichuris* and a positive serologic test for *Toxocara*, one subject had positive stool qPCR tests for *Trichuris* and *Giardia* as well as positive serologic testing for *Toxocara*, and the
third subject had positive serologic testing and stool qPCR for *S. stercoralis* and positive serologic testing for *Toxocara*; g - p-value=0.07, sensitivity of O&P compared to qPCR 12.5%, specificity 100%, positive predictive value (PPV) 100%, negative predictive value (NPV) 93.9%, kappa 0.21.

### Outcome assessment

With the exception of constitutional symptoms, which were slightly more likely to be present in subjects diagnosed with *Giardia* (p=0.01), there were no significant differences in any of the evaluated symptoms, exposures, or laboratory results between those found to be mono-infected with each of the different parasite species detected (e.g. *Toxocara* spp., soil-transmitted helminths, or *Giardia).* Consequently, for our primary analysis we compared uninfected individuals to subjects with evidence for any pathogenic parasitic infection, with infected subjects treated as a single group.

A large number of subjects (10/111 subjects who provided a sample for O&P testing, 9%) were found to have *Blastocystis hominis*. Although the pathogenicity of *B. hominis* remains controversial,[64, 65] infection with this organism in our sample was not associated with any significant differences in symptoms or laboratory values compared to the uninfected group. Accordingly, in our analysis *Blastocystis* was not included as a pathogenic parasitic infection.

### Demographic characteristics of subjects with parasitic infections

Nearly all of the subjects found to have evidence of a pathogenic parasite (14/15, 93.3%) had lived in the continental US for 2 years or less (compared to 70/110 of those who were uninfected, 63.6%, PR 6.8, p=0.02, Table 3). This history was therefore a very sensitive, though not specific, marker for parasitic infections (sensitivity 93.3%, specificity 36.4%, PPV 16.7%, NPV 97.6%). No other demographic variables, including age, gender, recruitment site, or region of origin differed significantly between those with and without infections. As shown in Table 3, there was a trend (though not statistically significant) towards a greater likelihood of parasitic infections in subjects of Asian origin, subjects who had not completed high school, and subjects with an annual household income of less than $20,000 per year.

**Table 3.** Findings associated with the presence of a parasitic infection

| Demographics | Uninfected<br>N=110 | Parasitic<br>Infection<sup>a</sup><br>N=15 | Prevalence<br>ratio<sup>b</sup><br>(95%<br>Confidence<br>Interval) | p-value |
| --- | --- | --- | --- | --- |
| **Length of time living in the**<br>**United States, N (%)** |  |  |  |  |
| > 2 years | 40 (36.4) | 1 (6.7) | Reference | **0.02** |
| ≤ 2 years | 70 (63.6) | 14 (93.3) | 6.8 (0.9-50.2) |  |
| **Mean age (standard**<br>**deviation)** | 31 (16.4) | 37.5 (21.6) | 1.0 (1.0-1.1) | 0.2 |
| **Gender, N (%)** |  |  |  |  |
| Female | 58 (52.7) | 8 (53.3) | Reference | 0.99 |
| Male | 52 (47.3) | 7 (46.7) | 1.0 (0.4-2.5) |  |
| **Country/Region of origin, N (%)** |  |  |  |  |
| Mexico | 26 (23.6) | 2 (13.3) | Reference | 0.28 |
| Asia (excluding India) | 24 (21.8) | 8 (53.3) | 3.5 (0.8-15.1) |  |
| India | 21 (19.1) | 2 (13.3) | 1.2 (0.2-8) |  |
| Central/South America | 18 (16.4) | 2 (13.3) | 1.4 (0.2-9.1) |  |
| Africa | 12 (10.9) | 1 (6.7) | 1.1 (0.1-10.8) |  |
| Middle East | 5 (4.5) | 0 (0) | Undefined |  |
| USA (Puerto Rico) | 4 (3.6) | 0 (0) | Undefined |  |
| **Years of schooling** |  |  |  |  |
| >12 | 56 (51.9)<sup>c</sup> | 5 (33.3) | Reference | 0.27 |
| 9-12 | 28 (25.9) | 4 (26.7) | 1.5 (0.4-5.7) |  |
| ≤8 | 24 (22.2) | 6 (40) | 2.4 (0.7-7.9) |  |
| **Annual household income, US dollars** |  |  |  |  |
| ≥20,000 | 34 (30.9) | 2 (13.3) | Reference | 0.2 |
| <20,000 | 76 (69.1) | 13 (86.7) | 2.6 (0.6-11.1) |  |

| Childhood Exposures (in country of origin) | Uninfected, number yes, (%)<br>N=110 | Parasitic infection, number yes, (%)<br>N=15 | Prevalence ratio<sup>b</sup> (95% confidence interval) | p-value |
| --- | --- | --- | --- | --- |
| Self-reported history of seeing worms in the stool | 18 (16.4) | 8 (53.3) | 4.4 (1.7-10.9) | **0.003** |
| Used a well as source of drinking water | 49 (44.5) | 11 (73.3) | 3.0 (1.0-8.9) | 0.05 |
| Raised in rural environment | 19 (17.3) | 3 (20) | 1.2 (0.4-3.8) | 0.73 |
| Frequently walked barefoot | 60 (54.5) | 11 (73.3) | 2.1 (0.7-6.2) | 0.27 |
| Close contact with animals | 51 (46.4) | 7 (46.7) | 1.0 (0.4-2.6) | 0.99 |
| Bathed in streams or ponds | 47 (42.7) | 7 (46.7) | 1.2 (0.4-3.0) | 0.79 |
| Lived in a house with a thatched roof | 20 (18.2) | 3 (20) | 1.1 (0.3-3.6) | 0.99 |
| Treated with antiparasitic medication (prior to emigration) | 38 (34.5) | 7 (46.7) | 1.6 (0.6-4.0) | 0.4 |
| Symptoms reported by the patient on the day of study enrollment<sup>d</sup> | Uninfected, number yes, (%)<br>N=110 | Parasitic infection, number yes, (%) | Prevalence ratio<sup>b</sup> (95% confidence interval) | p-value |

|  |  | N=15 |  |  |
| --- | --- | --- | --- | --- |
| Gastrointestinal | 26 (23.6) | 3 (20) | 0.8 (0.3-2.7) | 0.99 |
| Constitutional | 17 (15.5) | 6 (40) | 3 (1.2-7.5) | **0.03** |
| Musculoskeletal | 18 (16.4) | 6 (40) | 2.8 (1.1-7.1) | **0.04** |
| Dermatologic | 11 (10) | 1 (6.7) | 0.7 (0.1-4.7) | 0.99 |
| Pulmonary | 10 (9.1) | 1 (6.7) | 0.7 (0.1-5.1) | 0.99 |
| Cardiovascular | 7 (6.4) | 2 (13.3) | 2 (0.5-7.5) | 0.29 |
| Allergic | 2 (1.8) | 0 (0) | Undefined | 0.99 |

N – number of subjects; a – Not including non-pathogenic or disputed pathogenicity organisms; b – prevalence ratio comparing the frequency of the listed demographic variable, symptom, or exposure in those with versus those without evidence of a parasitic infection; c – 108 of the 110 subjects without parasitic infections responded to this question; d - gastrointestinal symptoms included: heartburn, difficulty or pain with swallowing, abdominal pain, diarrhea, nausea/vomiting; constitutional symptoms included: weight loss, fatigue, fevers, and weakness; musculoskeletal symptoms included pain and stiffness in the joints or myalgias; dermatologic: itching, hives, or rash; pulmonary symptoms: wheezing, shortness of breath, cough; cardiovascular: chest pain, palpitations, or irregular heart beat; and allergic symptoms included: those who responded “yes” to the question “do you currently have any symptoms of seasonal allergies.”

### Exposure history and symptoms in subjects with evidence of parasitic infection

A self-reported history of ever having seen worms in the stool was strongly associated with evidence of current or prior parasitic infection (8/15, 53.3% in the infected group reported this history versus 18/110, 16.4%, in the uninfected group, p=0.003, PR 4.4, Table 3). This history appeared to be a nonspecific marker of risk for any parasitic infection and was equally common among those diagnosed with enteric parasites compared to those with non-enteric parasites (p-value=0.8). Although it did not reach significance, there also appeared to be a trend towards increased parasitic infections in subjects who recalled drinking well water in their country of origin (11/15, 73.3%) compared to those who did not (49/110, 44.5%, p=0.05, PR 3). Other classic risk factors for parasitic infections (close contact with animals, *etc.)*, although commonly reported among study participants, did not differ between those with and without evidence of parasitic infections (all p-values >0.05, Table 3).

In the year prior to study enrollment, subjects with evidence of parasitic infections had experienced an overall increased number of physical complaints (median 8) compared to those with no evidence of parasites (4, p=0.02). However, symptoms typically thought of as indicative of parasitic infections, such as rash, pruritus, or diarrhea, did not differ significantly between the groups (all p-values >0.05 for symptoms experienced both the day of study enrollment and within the previous year).

### Laboratory values in subjects with evidence of parasitic infections

Both the median IgE level (249 IU/mL) and the proportion of subjects with an increased IgE level (7 of 15 subjects, 46.7%) were significantly increased in subjects with evidence for parasitic infections compared to those who were uninfected (median IgE 52 IU/mL, 22/110, 20%, subjects with elevated IgE level, both p-values <0.05, Table 4). In contrast, subjects who reported a history of seasonal allergies and/or asthma were not found to have increases in IgE (p-value=NS). The difference in median IgE levels between groups remained statistically significant when subjects <18 years old were excluded from analysis (as normal values for IgE are higher in this group). Although an elevated IgE level was a relatively specific finding for parasitic infections (specificity 80%, NPV 92%), this finding was not a sensitive marker of infection (sensitivity 47%, PPV 24%).

**Table 4.**
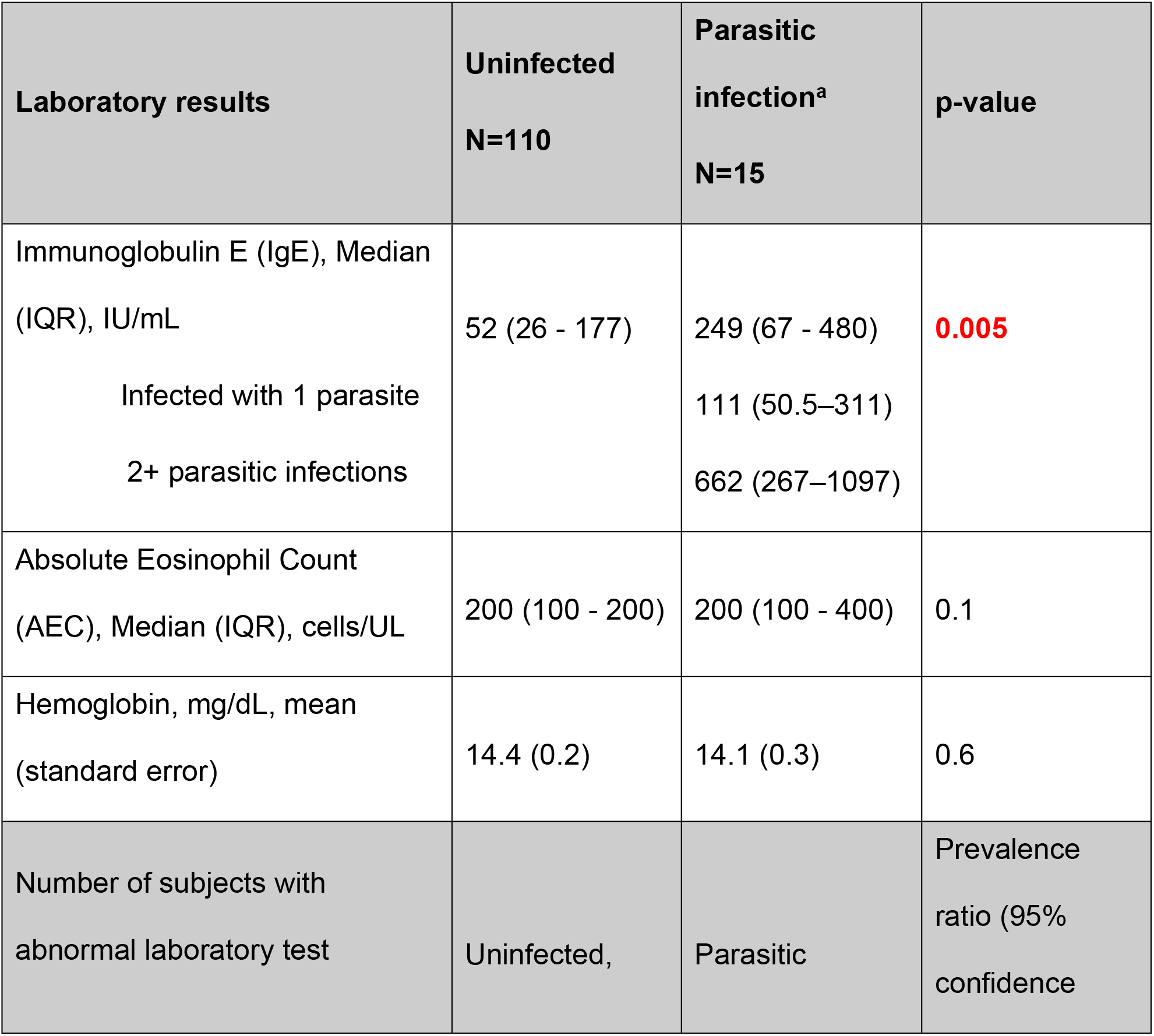
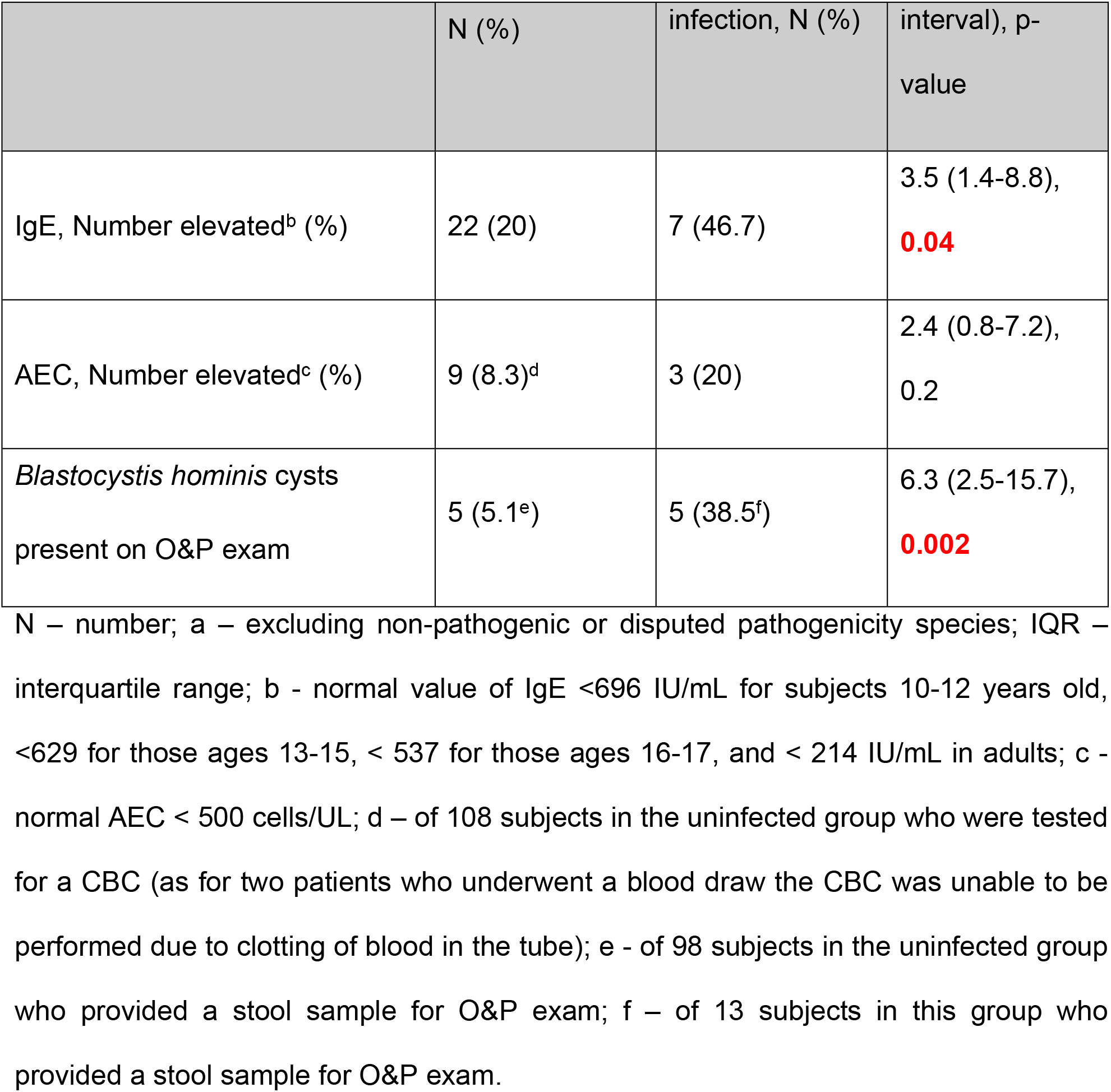
Laboratory results in uninfected versus infected individuals

The median AEC did not differ between those with and without evidence of a parasitic infection (Table 4). Eosinophilia in our sample was also not associated with a reported history of allergies, asthma, or taking medications known to cause eosinophilia.[66, 67] Despite a low sensitivity for parasites (20%, PPV 25%), however, eosinophilia was relatively specific to subjects with evidence of parasitic infection (specificity 91.8%, NPV 89.4%). The sensitivity of AEC as a marker for parasitic infection did not improve appreciably when only parasitic infections known to cause chronic eosinophilia *(Strongyloides* and Toxocara[28, 67–69]) were compared to those who were uninfected.

The presence of cysts of *B. hominis* on O&P exam was strongly associated with infection with a pathogenic parasite species. Five of the 13 subjects in the parasite positive group who returned a stool sample had a positive O&P for *Blastocystis*, 38.5%, compared to 5/98 tested for O&P exam in the uninfected group, 5.1%, p=0.002, PR=6.3.

## DISCUSSION

Despite the fact that many US immigrants come from areas where parasitic infections are endemic, data about the prevalence and risks for parasites among immigrants are sparse. In response to this deficit, we conducted a comprehensive evaluation for the presence of a broad array of parasitic infections in recent immigrants in Chicago. We found that 12 percent of recent immigrants in our sample had evidence of prior or current infection with a pathogenic parasite species.

Our study found a few key factors which were significantly more likely in subjects with evidence of parasites: a self-reported history of having previously observed worms in the stool, having immigrated within the past 2 years, and reporting a relatively high number of nonspecific physical ailments within the previous year. Our results also highlight that clinical risk factors (such as contact with animals and frequently walking barefoot), symptoms (including urticaria, rash, and diarrhea), and laboratory findings (eosinophilia) traditionally thought to be associated with parasites were commonly found but not predictive of infection in this study population. If confirmed in a larger sample, these results suggest that practitioners should have a high degree of suspicion for parasitic infections when evaluating recent immigrants regardless of the presence or absence of symptoms or laboratory abnormalities.

Our results highlight the limitations of current diagnostic tests for parasitic infections. For example, the sensitivity of O&P exam compared to qPCR testing (which is not yet universally available) for pathogenic parasites was very poor. While this could represent false positive qPCR tests, previous use of these tests in other studies has indicated a low rate of false positives.[8, 49, 61, 62] Further, the O&P exam has been shown to have serious limitations, in part because some parasitic infections are characterized by intermittent shedding in the stool.[70] The lack of adequate diagnostic tests for parasitic infections was also evident in the fact that, for many of the parasites screened for in this study, the primary method of diagnosis is serologic testing. Serologic tests can have significant limitations as they often have cross-reactivity among multiple parasites and these tests cannot distinguish between prior and current infections.[3, 6, 8, 17, 28, 37, 71, 72]

Similar to prior studies of immigrant populations,[28, 68] we found that a relatively high percent of subjects in our study had eosinophilia and/or elevated IgE levels. IgE appeared to be a slightly better marker for chronic parasitic infection in this sample than eosinophilia. However, neither test had an adequate sensitivity to merit recommending its use for routine screening purposes. For some parasitic infections, eosinophilia has been shown to be absent or mild with longstanding infection,[73, 74] and therefore the poor sensitivity of AEC for parasitic infections in our sample could reflect a lack of acute parasitic infections among these subjects. Although eosinophilia has long been thought of as a hallmark of helminth infection, several previous studies have also failed to show any correlation between the two.[75, 76] However, the high specificity of eosinophilia for parasitosis seen in our study suggests that, when this finding is present in a recent immigrant, a workup for parasitic infections should be initiated.

A large number of subjects in our study demonstrated cysts of *Blastocystis hominis* on O&P exam. The association between *Blastocystis* and clinical disease has been controversial. Various studies have shown that subjects with *B. hominis* may have an increased likelihood of irritable bowel syndrome and that symptomatic patients with this parasite may experience a beneficial effect from treatment with antiparasitic medications.[64, 77] In our study, however, infection with *Blastocystis* did not appear to be associated with any increase in symptoms in the preceding year or on the day of enrollment. Interestingly, the presence of *Blastocystis* cysts on an O&P exam was strongly associated with the presence of positive tests for pathogenic parasitic infections. Therefore, our results agree with those of several recent studies and suggest that the primary import of *B. hominis* infection may be from a public health perspective by serving as a surrogate marker for the presence of fecal-oral transmission.[65, 77, 78]

Our study results showed that a high percentage of recent immigrants had evidence of current or prior infection with a few key pathogenic parasites, including *Strongyloides, Toxocara*, and *Giardia.* Positive serologic testing for *Strongyloides* has been shown to be associated with a high likelihood of active ongoing infection and a risk for developing disseminated disease or hyperinfection syndrome.[17, 79] The rate of seropositivity for strongyloidiasis in our sample was 4%. This may have clinical implications because a study evaluating the cost-effectiveness of empiric treatment versus a test-and-treat strategy found that, if the prevalence of *Strongyloides* infection is greater than 2% in a community, then the presumptive treatment strategy is more cost-effective.[80]

Our study had several limitations. The small number of subjects found to have parasitic infections limited our ability to identify risk-factors associated with these infections. However, our sample size was necessitated by the extensive testing performed on each subject and was also similar to those of previous studies of immigrant health.[8, 81] Furthermore, the fact that our sample was very heterogeneous and subjects enrolled in the study came from many different countries could have reduced our ability to find associations between clinical factors and the presence of infection. Our recruitment of some subjects from English as a Second Language classes and at the Mexican consulate in Chicago could have skewed the study towards immigrants from non-English-speaking countries and Mexico. A significant limitation to this study, as for all studies evaluating parasitic infections (as well as to care of patients at risk for parasitic infection), is the lack of adequate diagnostic tests for parasitosis.

Despite these limitations, however, our study is one of the only recent studies to evaluate the prevalence of parasites in immigrants and provides a strong argument for further investigation of the health impacts of these infections in the immigrant community. Although our sample size was limited, a strength of our study is that we conducted a comprehensive evaluation for parasitic infections including testing of both stool and serum samples. Our results suggest that as many as 12% of recent immigrants in the community may have evidence of current or previous infection with a pathogenic parasitic species. From a public health perspective, it is important to note that the infections identified in the current patient sample are not typically spread from person to person. However, if confirmed in a larger study, these results present an important health disparity among a vulnerable underserved population in the US. This health disparity has persisted despite the presence of effective, safe, and well-tolerated antiparasitic medications capable of treating each of the identified pathogens in our sample.[17, 80]

## Acknowledgements

The authors would like to thank E. Scott Elder BS, Isabel McAuliffe PhD, and Sukwan Handali MD from the Centers for Disease control who performed the serologic assays conducted on patient samples. JH would like to thank Ed Mitre, MD, from USUHS and Thomas Nutman MD and Amy Klion MD from the NIH for input and help with the study design process. We also would like to thank the free clinics and social services organizations for collaborating with us and providing help with study recruitment. Dr. Herrick’s research was supported by the National Center for Advancing Translational Sciences, National Institutes of Health, through Grant UL1TR000050. Research funding support for RM was provided by the U.S. Department of Health and Human Services, Health Resources and Services Administration for Baylor College of Medicine Center of Excellence in Health Equity, Training, and Research (Grant No: D34HP31024). RM has also received some funding support from Romark Laboratory. The content is solely the responsibility of the authors and does not necessarily represent the official views of the NIH. The funders had no role in the study design, data collection and analysis, decision to publish, or preparation of the manuscript.

## Author contributions

Jesica A. Herrick was responsible for the funding acquisition, conceptualization of the study, study design and methodology, oversight of subject enrollment and regulatory aspects of the study, data curation and formal analysis, and writing of the manuscript. Monica Nordstrom, Patrick Maloney, and Miguel Rodriguez were involved in the investigation and recruited and enrolled subjects in the study and provided a critical review of the manuscript. Patrick Maloney was also responsible for the formal statistical analyses. Kevin Naceanceno and Gloria Gallo Enamorado performed the Multi-parallel real-time quantitative PCR (qPCR) conducted on patient stool samples as part of the investigation. Rojelio Mejia oversaw the qPCR conducted on patient stool samples (supervisory role) and provided a critical review of the manuscript. Ron Hershow was involved in a supervisory capacity in the study concept and design, analysis and interpretation of data, and writing of the manuscript.

## Supporting Information

**S1 Supplemental Appendix. Closed-answer questionnaire**. This questionnaire was filled out by each participant to obtain information about their demographics, symptoms, and exposure histories.

**S2 Supplemental Table 1. Serologic testing performed based on subjects’ countries of origin.**

**S3 Supplemental Figure 1. Study recruitment and enrollment.**

**S4 Supplemental Table 2. Responses to closed-answer questionnaire and laboratory results.** This table contains the complete individual responses subjects provided to the symptom and demographic questionnaire as well as the raw results from all laboratory tests performed.

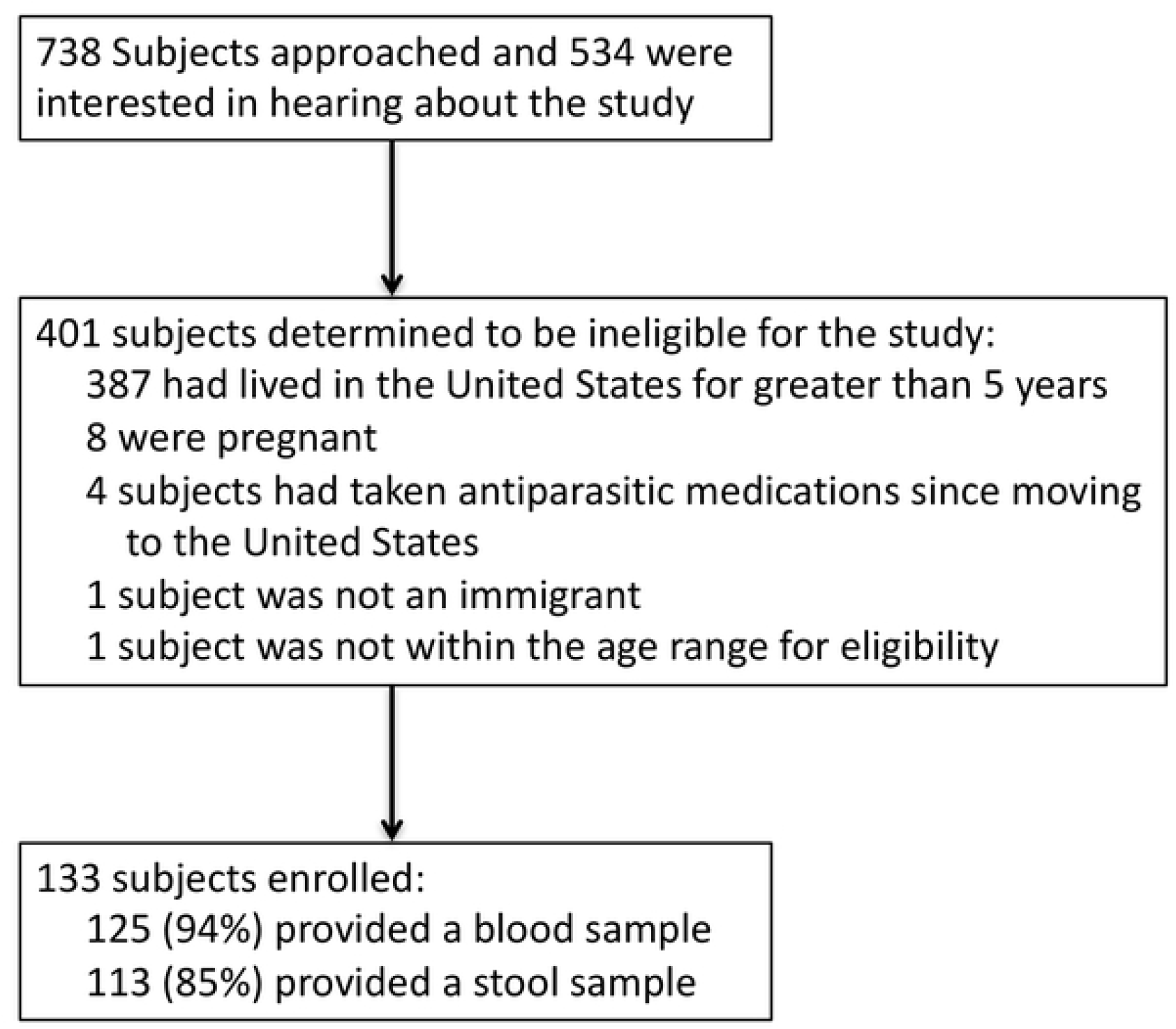

